# Single-cell anatomical analysis of human hippocampus and entorhinal cortex uncovers early-stage molecular pathology in Alzheimer’s disease

**DOI:** 10.1101/2021.07.01.450715

**Authors:** Jose Davila-Velderrain, Hansruedi Mathys, Shahin Mohammadi, Brad Ruzicka, Xueqiao Jiang, Ayesha Ng, David A. Bennett, Li-Huei Tsai, Manolis Kellis

## Abstract

The human hippocampal formation plays a central role in Alzheimer’s disease (AD) progression, cognitive traits, and the onset of dementia; yet its molecular states in AD remain uncharacterized. Here, we report a comprehensive single-cell transcriptomic dissection of the human hippocampus and entorhinal cortex across 489,558 cells from 65 individuals with varying stages of AD pathology. We transcriptionally characterize major brain cell types and neuronal classes, including 17 glutamatergic and 8 GABAergic neuron subpopulations. Combining evidence from human and mouse tissue-microdissection, neuronal cell isolation and spatial transcriptomics, we show that single-cell expression patterns capture fine-resolution neuronal anatomical topography. By stratifying subjects into early and late pathology groups, we uncover stage-dependent and cell-type specific transcriptional modules altered during AD progression. These include early-stage cell-type specific dysregulation of cellular and cholesterol metabolism, late-stage neuron-glia alterations in neurotransmission, and late-stage signatures of cellular stress, apoptosis, and DNA damage broadly shared across cell types. Late-stage signatures show signs of convergence in hippocampal and cortical cells, while early changes diverge; highlighting the relevance of characterizing molecular pathology across brain regions and AD progression. Finally, we characterize neuron subregion-specific responses to AD pathology and show that CA1 pyramidal neurons are the most transcriptionally altered while CA3 and dentate gyrus granule neurons the least. Our study provides a valuable resource to extend cell type-specific studies of AD to clinically relevant brain regions affected early by pathology in disease progression.

## Introduction

Late-onset Alzheimer’s disease (AD) is a progressive disorder characterized by stereotypical, decades-long accumulation of extracellular neuritic plaques and intracellular neurofibrillary tangles across multiple brain regions, a complex pathophysiological process commonly leading to clinical symptoms of cognitive impairment^1–4^. Great progress has been made in identifying the association of environmental and genetic risk factors for the disease, yet exploiting this knowledge for rational preventive and therapeutic intervention remains a major biomedical challenge^2,5^. Empirical case-control studies have analyzed the molecular composition of human brain tissue using genomic, transcriptomic, and epigenomic profiling technologies^6–10^. Diverse molecular pathways have been recurrently found to be dysregulated in AD, with alterations in metabolic, bioenergetic, proteostatic, cytoskeletal, inflammatory, and synaptic processes becoming broadly recognized as signatures of molecular AD pathophysiology^11,12^. The diversity of implicated processes suggests that both constitutive and specific homeostatic functions of interacting neuronal and glial cells are impaired in AD, and that distinguishing their relative contribution requires going beyond tissue-level resolution experiments^3^.

We recently reported the first single-cell transcriptomic analysis of AD, investigating gene expression alterations in cells of the prefrontal cortex, a brain region severely affected late in disease progression^13^. In addition to recognized AD-associated molecular pathways, our previous study showed prominent alteration of oligodendrocyte lineage cells, broad and cell-type-specific transcriptional dysregulation, and changes in the directionality of gene perturbations across cell types. Together, our previous observations highlighted the importance of single-cell-resolution studies for brain disease research, an approach now being applied to diverse human neurodegenerative^14–16^, and neurodevelopmental disorders^17,18^.

The hippocampus (HIP) is one of the earliest brain structures affected in AD progression, and it is also a central locus for several domains of cognitive function altered in AD^19^. Despite its central importance in dementia and neurodegeneration, however, the human hippocampus and surrounding retrohippocampal regions remain uncharacterized at single-cell resolution in the context of aging and AD pathology. The hippocampus is part of the hippocampal formation, a brain structure additionally composed by the medial and lateral entorhinal cortex (EC), the subicular complex (Sub), and the dentate gyrus (DG)^19^. These structures display early progressive signs of pathology and are known to host selectively vulnerable neuronal populations ^20–22^. Only a handful of studies have analyzed hippocampal transcriptional profiles of single-cells in mice^23–25^ or pathology-free human samples^26,27^. Characterizing the transcriptional architecture of the human hippocampal region at single-cell resolution in the context of aging and pathology is crucial for understanding the molecular pathophysiology of AD onset and progression across glia, neurons, and their functional circuits.

Here, we report a comprehensive single-cell dissection of the aged human hippocampus and entorhinal cortex, enabling the molecular characterization of their cellular diversity and the transcriptional manifestation of AD in early-affected structures. We profile a total of 112 human post-mortem brain samples from 65 aged individual donors with variable degrees of pathology, generating and analyzing 489,558 high-quality single-nucleus transcriptomes. We integrate our data with existing human and mouse tissue-microdissection, neuronal sorting, spatial, and single-cell transcriptomic datasets to aid interpretation and annotation of the human cell diversity uncovered. Our results reveal a unique cellular-level view of neuropathology in the human hippocampus and entorhinal cortex, demonstrate that human cellular expression patterns capture fine-resolution neuronal anatomical topography, and reveal modular gene expression alterations that manifest early or late in pathology progression across brain structures and cell types. Overall, our analysis reveals the importance of sampling early-affected brain regions and early pathological stages to characterize progressive patterns of molecular pathology. Our generated data, cell annotations, and expression patterns provide a valuable resource for investigating early Alzheimer’s disease pathology at single-cell resolution.

## Results

### Single-nucleus RNA-seq profiling of human hippocampus

We selected 31 individuals with positive pathological diagnosis of AD (AD group) and 34 individuals with negative diagnosis (no AD pathology group, NoAD), balancing for male:female ratio (16:15 in AD, 17:17 in NoAD), age (72-95 years, average 85:84), and postmortem interval (0.86-12 hours, average 6:6.5). Considering pathology staging, we selected a balanced number of subjects with early/limbic (Braak stage 3 or 4, n=17); and late/neocortical (Braak 5 or 6, n=17) involvement of neurofibrillary tangle (NFT) pathology (**Fig. 1a**). AD subjects present high levels of both β-amyloid and Tau pathology, as well as increased cognitive decline; with incremental levels of both pathology types from early to late Braak stages, and with no differences in age or post-mortem time intervals (pmi) (**Extended Data Fig. 1a, b**). All individual donors are participants in the Religious Order Study (ROS) or the Rush Memory and Aging Project (MAP), collectively known as ROSMAP^28^. From 43 of the individuals, we obtained postmortem tissue samples extracted from both the hippocampus and the entorhinal region, from 19 only hippocampal, and from 3 only entorhinal samples. Therefore, we processed a total of 62 hippocampal and 48 entorhinal samples encompassing 65 different individuals. All tissue samples were obtained without prior microdissection of anatomical subdivisions (**Fig. 1b**). To characterize the cellular transcriptomic landscape of the samples we performed single-nucleus RNA sequencing (snRNA-seq) using 10x Genomics Chromium chemistry v3 platform, producing a dataset of 574,582 single-nucleus transcriptomes. We processed and filtered the initial data based on standard quality metrics (Methods), resulting in 489,558 high-quality cells (85%) that we analyzed for the rest of the manuscript. To this end, we designed an integrative computational framework combining graph-clustering algorithms to identify coherent cellular populations, with external human and mouse data sources to facilitate interpretation of transcriptomic patterns (**Fig. 1b**). Data generated herein will be publicly accessible with a data use agreement through the ROSMAP data compendium (see ‘Data availability’).

**Figure 1.**
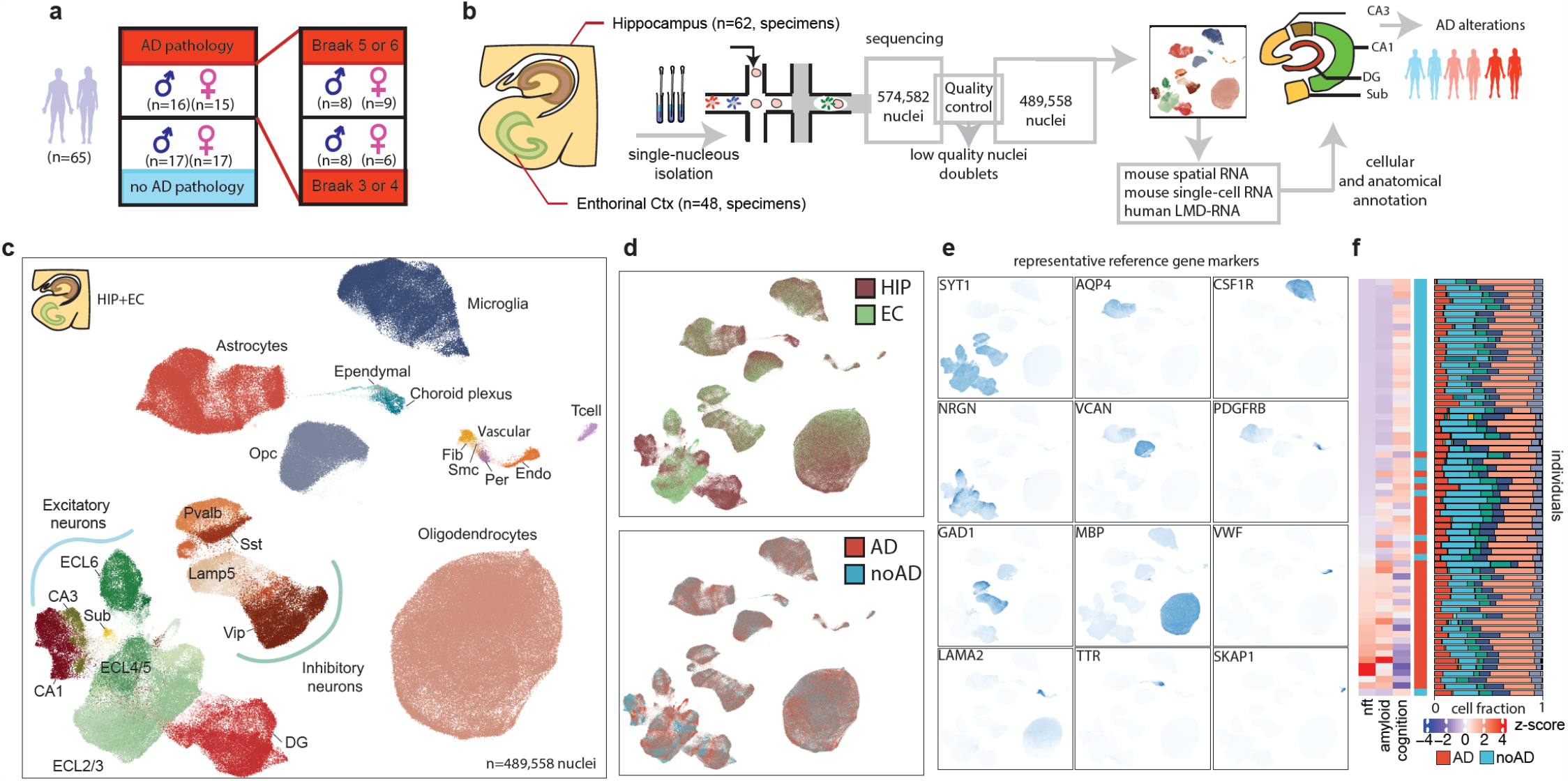
snRNA-seq profiling and cell type annotation. **a**, Study cohort description by AD pathology groups: positive pathological diagnosis (red), negative diagnosis (blue). Pathology groups are further divided according to Braak stage. **b**, Experimental design couples RNA sequencing of nuclei isolated from hippocampal and entorhinal samples with computational analysis for stringent QC and extensive cellular and anatomical annotation to characterize AD-associated cell-type specific and stage-dependent transcriptional alterations. **c-d**, Two-dimensional ACTIONet representation of all high-quality cells (n=489,558) included in downstream analysis and labeled by major cell types and neuronal populations (**c**), by brain region (**d**, top), and by AD status (**d**, bottom). **e**, Expression patterns of representative marker genes projected across cells localize in cell type neighborhoods consistent with annotations. **f**, Neuropathological measurements and fraction of cells of a given type (columns) by individual donor (rows).

### Cellular diversity of human hippocampus and entorhinal cortex

To identify and annotate major cell types and subpopulations, we performed two rounds of graph-based clustering analysis (**Extended Data Fig. 1c, Methods)**. We used the cell groups resulting from the first round to identify major brain cell types. On the basis of expression patterns and statistical overrepresentation of known marker genes, we reproducibly identified cell clusters matching signatures of 8 major brain cell groups supported by independent reference sources^29,30^ (**Extended Data Fig. 1d,e**). Cell groups include excitatory (Ex) and inhibitory (In) neurons, astrocytes (Ast), oligodendrocytes (Oli), oligodendrocyte progenitor cells (Opc), microglia (Mic), vascular cells (enriched for endothelial and pericyte markers), and choroid plexus cells.

To further dissect cell heterogeneity at higher resolution, we performed within-cell group analysis in a second clustering round, identifying multiple subgroups for each major cell group. Similar subcluster marker-based annotation identified small coherent subgroups consistent with T-cell, Fibroblast (Fib), Endothelial (Endo), Pericyte (Per), Smooth muscle cell (SMC), and Ependymal cell signatures. By integrating annotations at both levels, we defined unified labels for 13 major cell types (**Fig. 1c, Extended Data Fig. 1d, e)**. Because neuronal cells displayed the most apparent within-cell type variability across regions (HIP, EC), and between AD diagnosis groups, as evidenced by segregating patterns in 2D projection plots (**Fig. 1c, d)**, we performed an independent analysis to characterize and interpret neuronal subpopulations more thoroughly (see subsection below). This analysis resulted in the identification of subpopulations from all major hippocampal subregions (DG, CA fields, Sub, EC), as well as cardinal interneuron subtypes (Sst, Vip, Pvalv) (**Fig. 1c**).

Genes preferentially expressed across cell types showed strong enrichment of reference gene marker sets^29,30^ (**Extended Data Fig. 1e)**, and cell type expression profiles were strongly correlated between donors, with higher correlation (average Pearson r=0.96) for high-abundance cell types (average number of cells = 1,229 per donor) and lower (r=0.78) for low-abundance cell types (average number of cells = 22 per donor) (**Extended Data Fig. 1f-g)**. To corroborate cell annotations, we confirmed that cross-cell expression patterns of well-established gene markers clearly define cell neighborhoods within a 2D projection plot. Neuronal cells are marked by SYT1 expression, which further separate into two major cell groups corresponding to excitatory (NRGN) and inhibitory (GAD1) neurons (**Fig 1e**). Glial cells separate oligodendrocyte (MBP), oligodendrocyte progenitor (VCAN), astrocytes (AQP4), and microglia (CSF1R) cells. In addition, low-abundance cell groups showed localized expression of markers of immune and vascular-related cells, including T-cells (SKAP1), endothelial cells (VWF), pericytes (PDGFRB), and fibroblasts (LAMA2, CEMP); as well as 2 small groups of ependymal and choroid plexus cells (TTR). Cell type proportions are broadly consistent across donors and pathological groups (**Fig. 1f**).

### Single-cell patterns recover hippocampal spatial cytoarchitecture

To further characterize neuronal diversity, we performed an independent analysis focusing only on neuronal cells. We computationally sorted neuronal nuclei from the HIP and EC datasets to build and cluster an integrative cell network for each neuronal class (Excitatory Glutamatergic and Inhibitory GABAergic neurons) (**Fig. 2a, b, Extended Data Fig. 2a, b**). After data integration, subsets of glutamatergic neurons clearly segregated by brain region, a pattern not observed in GABAergic neurons. The two neuronal classes show contrasting mixing patterns in 2D-projection plots (**Fig. 2c**), with only glutamatergic neurons having clusters constituted exclusively of HIP cells. To quantify this pattern, we estimated for each cluster the distribution of cells isolated from each region and confirmed a strong frequency imbalance between brain regions across clustering subgroups of glutamatergic but not GABAergic neurons (**Fig. 2d, e**). This result is consistent with observations from cortical studies in humans^13^ and mouse^31^, suggesting that, unlike other neuronal classes or glial cells, excitatory neurons show strong region specific transcriptional patterns.

**Figure 2.**
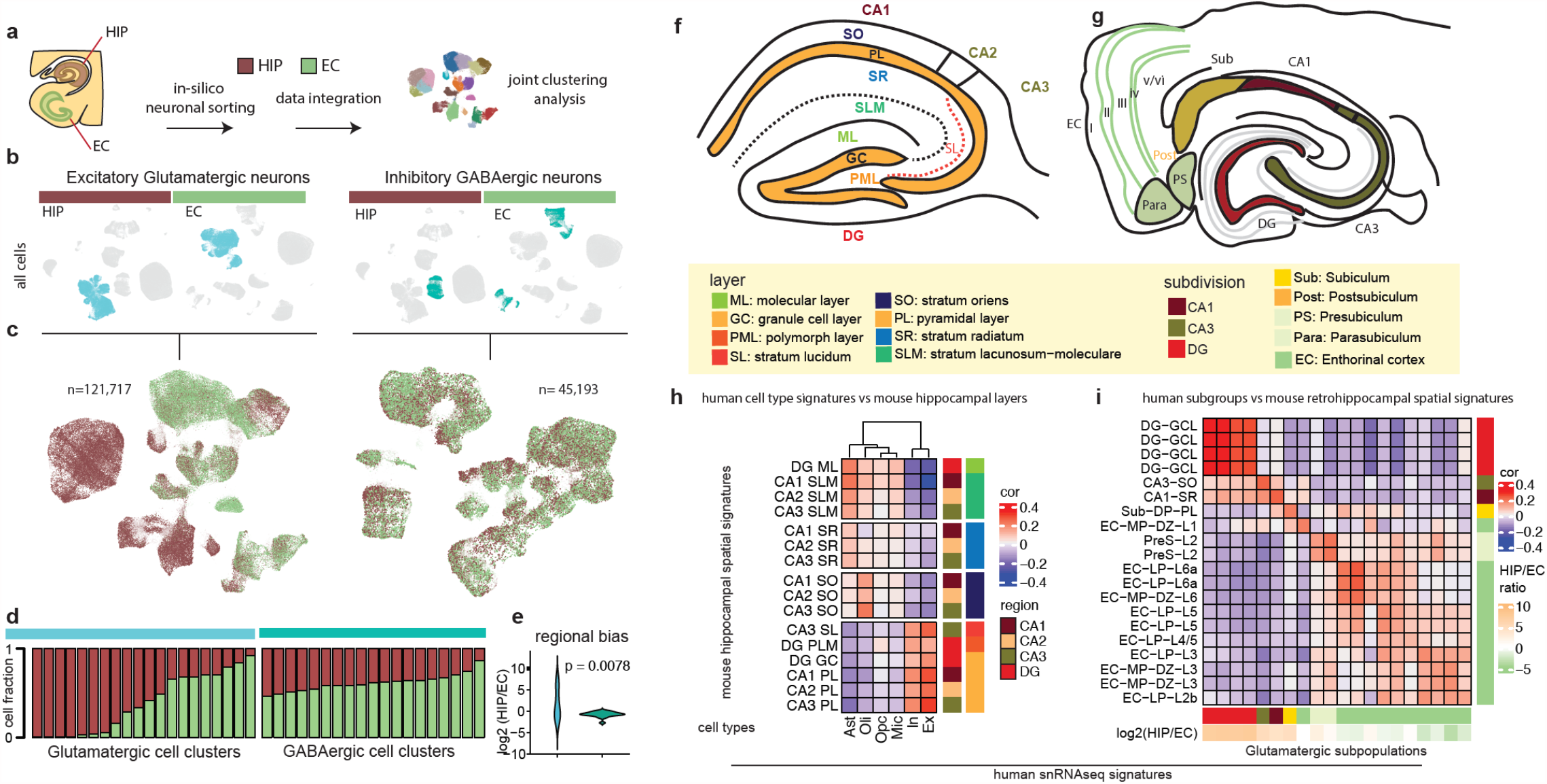
Integrative neuronal analysis and cytoarchitectural interpretation. **a**, Joint clustering analysis of HIP and EC cells used to identify neuronal subgroups. **b**, Projection of glutamatergic (blue) and GABAergic (green) neurons to cell space by brain region. **c**, Two-dimensional ACTIONet representation of glutamatergic (n=121,717) and GABAergic (n=45,193) neurons labeled by brain region (HIP, brown, EC, green). **d**., Fraction of glutamatergic (left) and GABAergic neurons isolated from HIP (brown) of EC (green) samples by subpopulations identified by joining clustering (columns). **e**, Comparison of the ratio of HIP to EC fractions recovered in glutamatergic (blue) vs GABAergic (green) neurons. P-value based on a two-sided wilcoxon rank-sum test. **f-g**, Schematic illustration of hippocampal layer cytoarchitecture (**f**) and major hippocampal formation subdivisions (**g**) modeled after mouse reference. **h**, Similarity between human cell type profiles and mouse hippocampal layered signatures computed from spatial transcriptomics data. **i**, Similarity between human glutamatergic subpopulation signatures and mouse hippocampal and retrohippocampal substructure profiles computed from spatial transcriptomics data. Similarity is measured using the pearson correlation coefficient.

To interpret the transcriptomic heterogeneity uncovered through joint clustering analysis, and to investigate its potential relationship with anatomical substructure and spatial cytoarchitecture, we compiled existing human and mouse transcriptomic data considering laser-microdissected^32^, spatial^33^, single-cell^24^, and neuron-sorted^34^ information. For each available dataset, annotated anatomical subregion, and neuron type, we quantified expression and signature profiles that we then used as reference to compare and interpret our single-nucleus data (Methods, **Supplementary Table 1**). We considered cell-sorted references for pyramidal neurons of the CA subfields and subiculum, granule cells of the dentate gyrus, and tissue references for all major subdivisions of the hippocampal formation (DG, CA fields, Sub, EC, and layers therein).

We first examined the relationship between cell type expression profiles and the anatomical distribution of principal cells in the hippocampus. Consistent with hippocampal cytoarchitecture (**Fig. 2f**), correlation analysis of human snRNA cell types (**Supplementary Table S2**) with mouse spatial transcriptomes (**Supplementary Table S1**) showed that cell type signatures robustly segregate samples by layer rather than subdivision, with presence or absence of neuronal signatures as the most discriminative pattern (**Fig. 2h**). The subdivisions of the hippocampus are structured in 4 and 3 layers spanning the CA fields (SLM, SR, PL, SO) and dentate gyrus (PL, GC, ML), respectively (described in **Fig. 2f, g**). In accordance with layer cytoarchitecture, PL and GC layers clustered together, showing high correlation with neuronal signatures; while SLM and ML correlated with glial signatures (**Fig. 2h, Extended Data Fig. 2c**). SO, and SR layers of the CA subfields showed a less defined pattern, presenting weaker correlations with oligodendrocyte and astrocyte cell signatures, respectively; as well as absence of neuronal transcriptional signatures (**Extended Data Fig. 2c)**. These patterns are consistent with observed cell body and commissural fiber distributions across subfields and layers^19^, explaining the clear segregation of neuronal and nonneuronal signatures.

The close correspondence between recovered snRNA cell type patterns and hippocampus cytoarchitecture indicates that transcriptional variation across hippocampal subdivisions strongly reflects the distributions of the major transcriptional cell classes uncovered with snRNA-seq and suggests that the mouse spatial transcriptomic atlas^33^ provides an appropriate comparative reference for human data interpretation. To investigate whether similar conclusions emerge from human tissue, we profiled a subset of subdivisions and layers from 2 pathologically-unaffected donors using laser capture microdissection followed by RNA sequencing (LCM-seq). We observed patterns highly consistent with those observed when comparing with mouse data (**Supplementary Fig. 2d**), further supporting the potential of snRNA-seq to dissect tissue-level heterogeneity and the suitability of cross-species analysis to aid interpretation.

### Molecular characterization of neuronal diversity

Given that layers and not anatomical subdivisions are primarily discriminated by the presence or absence of neuron signatures (**Fig. 2h**), we next tested whether the specificity of different human neuronal subpopulations recovered by joint clustering analysis (**Extended Data Fig. 2a, b**) recapitulates hippocampal and entorhinal substructure. We focused on excitatory glutamatergic neurons, which have a spatially and anatomically patterned cytoarchitecture^19^. Notably, when measuring the transcriptomic similarity between human subpopulations and mouse spatial segments, we found that human excitatory neurons capture distinctive expression patterns that broadly group together adjacent hippocampal formation subdivisions (**Fig.2i, Extended Data Fig. 3a**), suggesting the embedding of anatomical cytoarchitectural information within unbiased snRNA profiles. Correlation analysis and hierarchical clustering segregated 4 neuronal subpopulations strongly correlating with the granule cell layer of the dentate gyrus (DG), and 2 unique subgroups strongly correlating with the pyramidal layer of the CA1 and CA3 subfields. The remaining clusters mapped to areas spanning the subicular complex and different layers of the EC in the mouse (**Fig.2i, Extended Data Fig. 3a**). We leveraged this anatomical information to infer interpretable distinctions underlying the cluster partitioning patterns of human neuronal cells.

To further interpret the identity of human neuronal subpopulations, we extracted, reanalyzed, and compared mouse hippocampal and retrohippocampal single-cell transcriptomic data reported as part of the Allen Institute’s cell type database ^24^. Transcriptomic similarity between human subpopulations and mouse reference cell types confirmed most of the associations recovered by tissue-resolution spatial transcriptomic comparison, while suggesting higher resolution mapping identities for 5 subpopulations without a clear anatomical assignment (**Extended Data Fig. 3a, b**). To define a consensus interpretation and annotation for all human subpopulations, we next integrated the best-matching anatomical regions and cell types obtained from mouse spatial and single-cell transcriptomic comparisons with those obtained in analogous comparisons with mouse cell-sorted neurons and human microdissected tissue samples (**Extended Data Fig. 3c, Supplementary Table S3**). Applied to all excitatory clusters, this analysis resulted in the identification of 14 well-defined glutamatergic subpopulations, with signatures of DG granule cells (DG), and pyramidal neurons from subfields CA1 and CA3, entorhinal cortex (ECL2-6), Subiculum (Sub), para-, post-, and presubiculum (PPP); and 3 smaller groups. Best mouse cell type matches suggest that these latter groups represent cells from the area prostriata (APr), deep layer neurons marked by Car3 (Car3L6), and a heterogeneous group with mixed signatures reminiscent of Cajal–Retzius cells along with progenitor and neuronal markers (CRmx) (**Fig. 3a**). We confirmed that the identified subpopulations show similar relative distributions and associations at single-cell level in human and mouse, as evidenced by the spatial distribution in 2D projection plots (**Fig. 3a-c**), as well as similar preferential expression patterns of marker genes (**Fig. 3d, e**), and global transcriptional similarity (**Fig. 3f**).

**Figure 3.**
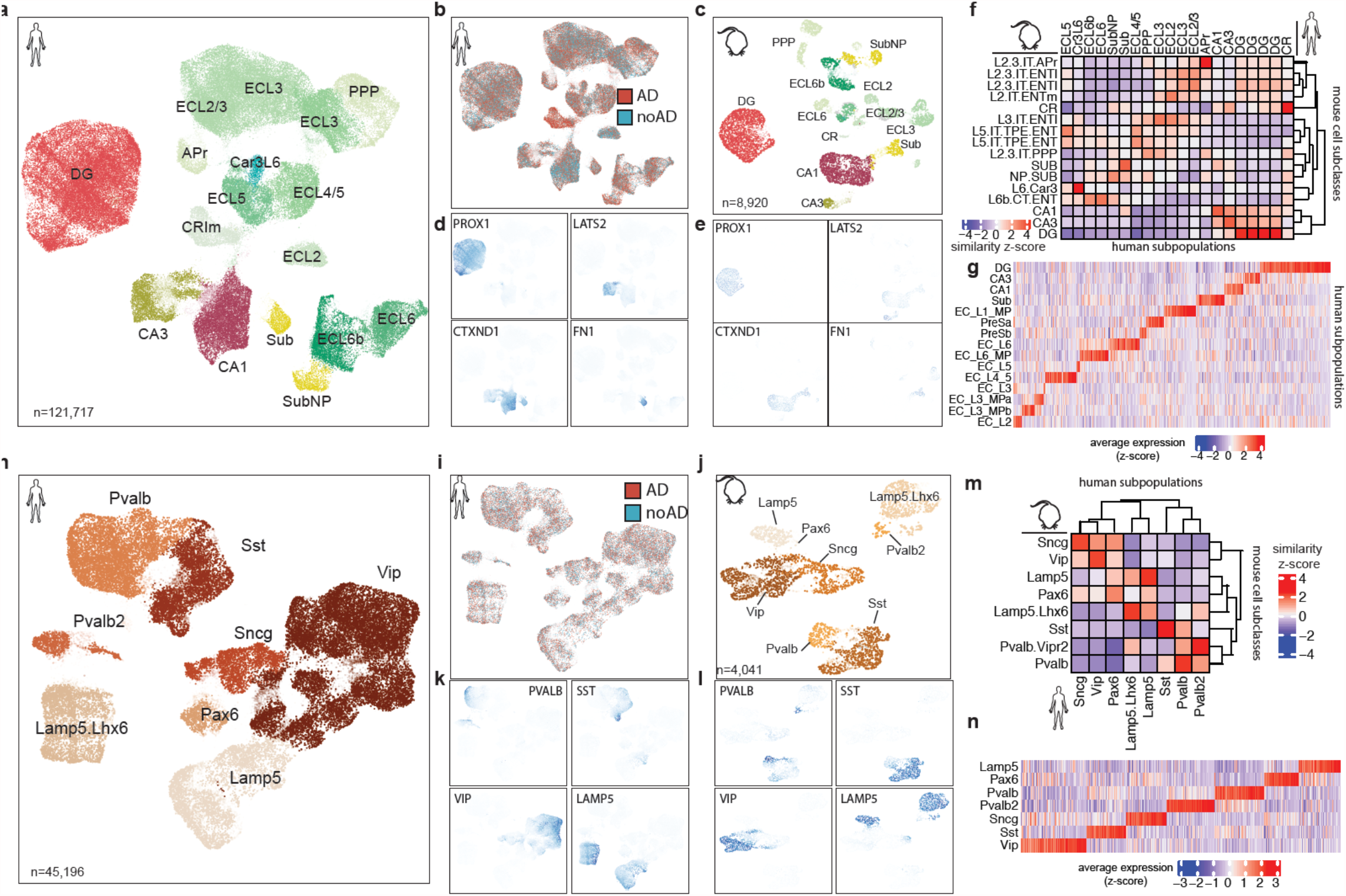
Annotation of neuronal subpopulations. **a-b**, Two-dimensional ACTIONet representation of glutamatergic neurons (n=121,717) labelled by subpopulations deduced from integrative data interpretation (**a**) and AD status (**b**). **c**, Two-dimensional ACTIONet representation of glutamatergic neurons reanalyzed from mouse (retro)hippocampal scRNA data reported by the Allen institute of Brain science and labeled by analogous subpopulation. **d-e**, Expression of representative orthologous glutamatergic marker genes projected across cells localize in subpopulation neighborhoods consistent with annotation in human (**d**) and mouse (**e**). **f**, Similarity between human glutamatergic subpopulation signatures (columns) and mouse hippocampal and retrohippocampal transcriptional cell types of the Allen Institute cell type reference (rows). **g**, Average expression patterns of genes detected as preferentially expressed across human glutamatergic subpopulations. **h-i**, Two-dimensional ACTIONet representation of GABAergic neurons (n=45,196) labelled by subpopulations deduced from comparison with reference mouse transcriptomic cell types (**h**) and AD status (**i**). **j**, Two-dimensional ACTIONet representation of GABAergic neurons reanalyzed from mouse (retro)hippocampal scRNA data reported by the Allen institute labeled by analogous subpopulation. **k-l**, Expression of representative orthologous GABAergic marker genes projected across cells localize in subpopulation neighborhoods consistent with annotation in human (**k**) and mouse (**l**). **m**, Similarity between human GABAergic subpopulation signatures (columns) and mouse transcriptional cell types of the Allen Institute cell type reference (rows). **g**, Average expression patterns of genes detected as preferentially expressed across human GABAergic subpopulations. Similarity between human and mouse signatures is measured using the Pearson correlation coefficient.

Unlike glutamatergic neurons, GABAergic neurons did not show strong regional segregation (**Fig. 2c, d**), and thus we interpreted GABAergic neurons based on similarity with reference GABAergic mouse cell types, rather than their anatomical structures (**Extended Data Fig. 3d**). We identified 5 human neuronal subpopulations consistent with signatures of major GABAergic subclasses (Pvalb, Pvalb2, Sst, Vip, Lamp), and 3 additional subpopulations mapping to mouse Lamp.Lh6, Pax6, and Sncg types. We confirmed that this level of interpretation recovers similar relative distribution and association of subpopulations at single-cell level in human and mouse (**Fig. 3h-j**), and consistent expression patterns of marker genes (**Fig. 3k, l**) and global transcriptional similarity (**Fig. 3m**). In addition to interrogating the expression of known marker genes, we identified discriminatory genes that show highly specific expression (Bonferroni-corrected pval<0.01, logFC>1, negative binomial mixed model (NBMM)) across human excitatory and inhibitory subpopulations (**Fig. 3g, n; Supplementary Table S4**).

### Modular AD-associated expression changes

We next used our high-resolution cellular and spatial annotations to study whether the molecular manifestation of AD pathology differs across regions, cell types, and neuronal subpopulations (**Fig. 4a**). To systematically characterize and dissect such differences at cell-type resolution, we measured AD-associated expression changes independently for each dissected brain region (HIP, EC) and for each major cell type, while correcting for age, sex, and postmortem time interval. In addition, because the AD phenotype is inherently complex and heterogeneous^13,35^, we considered AD effects at two neuropathological stages: changes happening at an early/limbic (Braak 3 or 4) or late/neocortical (Braak 5 or 6) stage, each relative to stages with low or undetectable pathology (Braak stage 0, 1, or 2) (**Supplementary Table S5**). We confirmed that our pathological group definitions were reflected in both β-amyloid and NFT pathology, which were both higher in late-stage individuals, and in cognition, which was lower in late-stage individuals (**Extended Data Fig. 1a, b**).

**Figure 4.**
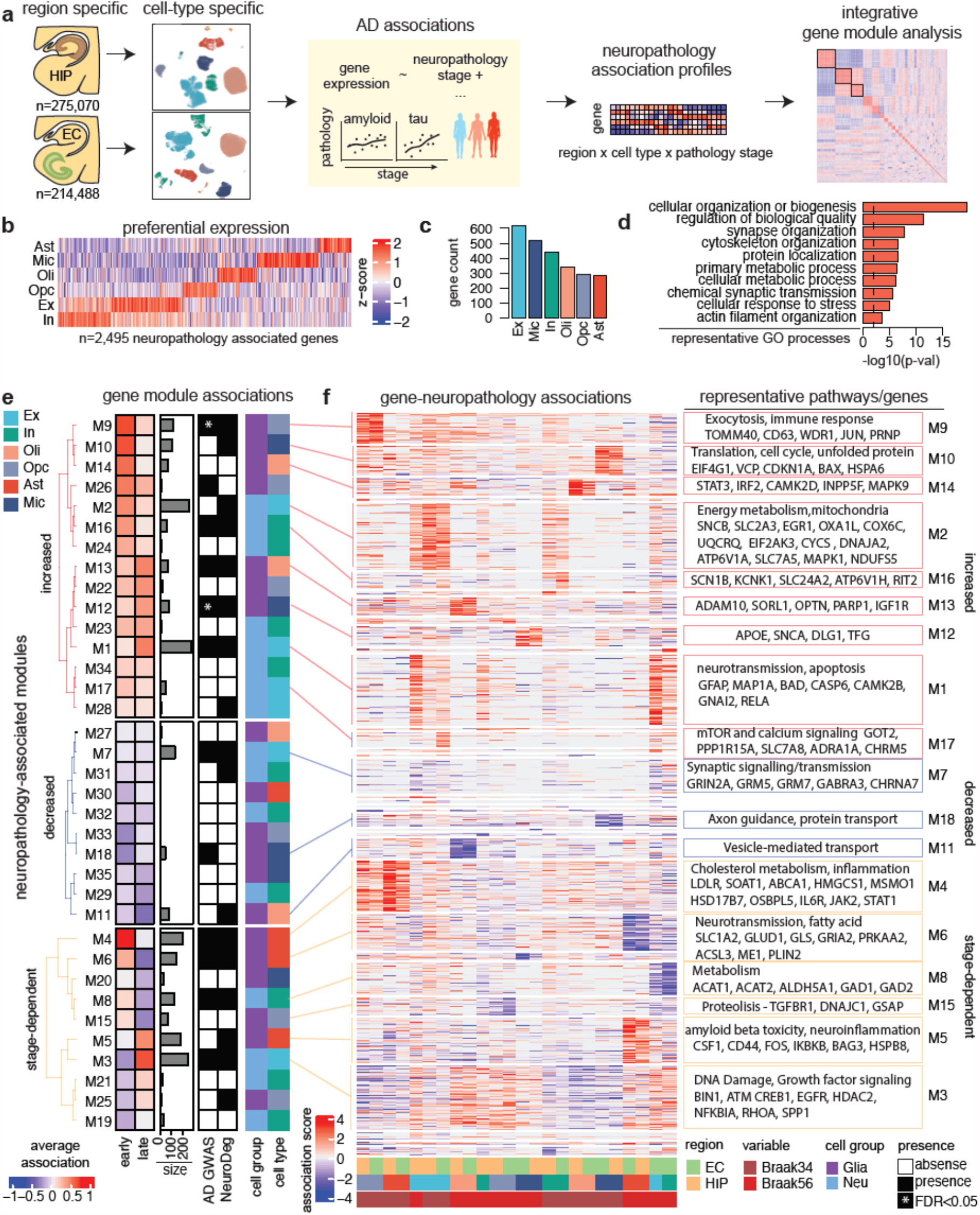
Modular stage-specific gene expression changes in AD pathology. **a**, Analysis pipeline consists of (I) the estimation AD-associated expression changes happening at an early/limbic (Braak 3 or 4) or late/neocortical (Braak 5 or 6) stage, relative to stages with low or undetectable pathology (Braak stage 0, 1, or 2), and independently by brain region (HIP, EC) and major cell type; (II) the definition of a neuropathology association profile by concatenating gene association scores for each combination of neuropathology stage, brain region, and cell type; and (III) the identification of gene modules based on neuropathology association profile similarity. **b**, Relative average expression of neuropathologically-associated genes across major cell types. **c**, Number of neuropathology associated genes with preferential expression in a given cell type (bars). **d**, Representative GO biological processes overrepresented within neuropathology associated genes. **e-f**, Modules of neuropathologically-associated genes capture stage-dependent and cell type specific changes as measured by average module association scores (**e**) and gene-neuropathology association scores (**f**). Association scores are computed by the negative logarithm of the p-values estimated by a negative binomial mixed model signed by the directionality of expression change.

We identified 2,495 neuropathology-associated genes that show at least one significant expression change (p-value < 0.01, negative binomial mixed model (NBMM)) across comparisons. Subsets of these genes are expressed preferentially in all major neuronal or glial cell types (**Fig. 4b**), with the largest subset being preferentially expressed in excitatory neurons, followed by microglia (**Fig. 4c**), suggesting that neuropathology is associated with functional alterations in multiple cell types. However, despite a dominant pattern of cell-type-preferential expression (i.e., genes having higher expression in a given cell type relative to others), we also observed that a large portion of neuropathology differential genes show broad base level expression (detected in >10% of the cells of a given type) in more than one cell type, with 31.5% (n=786) of the genes being expressed in all major cell types (**Extended Data Fig. 4a**). These broadly-expressed neuropathology-associated genes include ADAM10, SORL1, BIN1, PARP1, IGFR1, AKT3, and EIF2AK4, which indeed involve core cellular functions, including cellular metabolism, intracellular transport, and stress response (**Extended Data Fig. 4a, Supplementary Table S6**). GO enrichment analysis supports this observation, with an overrepresentation of genes involved in metabolic, bioenergetic, proteostatic, cytoskeletal, inflammatory, and synaptic biological processes within the complete set of neuropathology-associated genes, indicating that these broadly-acting biological processes are systematically altered across multiple cell types in AD (**Fig. 4d, Supplementary Table S6**).

To analyze the specific context in which expression alterations occur, we grouped all neuropathology-associated genes into modules, based on the similarity of their association scores for each combination of neuropathological stage, brain region, and cell type (together defining a gene-wise neuro-pathology association profile) (**see Methods, Supplementary Table S7**); and then characterized groups of genes with similar association profiles (**Fig. 4a, Supplementary Fig. S4b-c**). We identified 35 gene modules with consistent association patterns across stages, brain regions, and cell types; ranging from 5 to 272 genes in size (median=45 genes) and capturing both glial and neuronal alterations (**Fig. 4e, f; Supplementary Table S8**). Hierarchical clustering of gene association scores (**Fig. 4f**) primarily grouped pathological stages and cell types, rather than brain regions; suggesting that stage-dependent cell type association with pathology are broadly consistent between hippocampal and entorhinal cell types, with few exceptions in neurons. Notably, we identified three distinct categories of modules with respect to their stage-dependent response to pathology (**Fig. 4e**). Two groups were characterized by consistently increased (n=15 modules, 1,147 genes) or decreased (n=10 modules, 294 genes) expression levels across pathology stages and involve changes in all cell types, except astrocytes. The remaining group (n=10 modules, 1,034 genes) includes modules with a stage-dependent and qualitatively distinct response to pathology, presenting increased followed by decreased expression or vice versa.

Consistently-increased modules with strong early response in OPCs and oligodendrocytes (M9, M14) implicate exocytosis, immune response, and inflammation genes (e.g., TOMM40, CD63, STAT3, and IRF2); those with increased expression in microglia (M10) implicate cell cycle, protein translation, and unfolded protein response genes (e.g., EIF4G1, CDKN1A, and BAX); while early neuronal modules (M2, M16) implicate energy metabolism, mitochondria, and oxidative phosphorylation genes (e.g, EGR1, COX6C, OXA1L, ATP6V1H). Consistently-increased modules with a strong late response capture the upregulation of AD GWAS risk genes such as ADAM10, SORL1, PARP1, APOE, and SNCA in glial cells (M13, oligodendrocytes; M12, microglia); and of genes involved in neurotransmission and apoptosis in neurons (e.g., BAD, CASP6, CAMK2B, GNAI2). Consistently-decreased modules were less prominent and involve synaptic signaling, axon guidance, and protein transport genes; including major neurotransmitter receptors (e.g., GRIN2A, GRM5/7, GABRA3, CHRNA7) affected in neuronal cells (**Supplementary Fig. S4d)**.

Approximately half of the modules (n=19, ∼54%) include one or more genes previously associated with AD and/or with another major neurodegenerative disorder (PD, ALS, HD, dementia), including TOMM40, APP, SLC2A3, PARP1, APOE, and SNCA. A total of 19 AD-associated GWAS genes are also differentially expressed and are included within modules. Furthermore, 2 of the modules (M9 and M12) are significantly associated with gene-level GWAS risk scores for AD (FDR <0.05, ranked GSEA) with M9 having the strongest association (FDR <0.0005). In contrast, we did not find any association between AD neuropathology-associated modules and GWAS risk for either schizophrenia or type 2 diabetes (**Fig. 4e, f, Supplementary Table S8**), indicating that differentially expressed genes capture AD risk genes specifically. Notably, while module M12 includes the apolipoprotein APOE and is specifically perturbed in microglia, consistent with well-established associations between AD risk and cell type specificity^13,36^; the more strongly associated M12 includes the mitochondrial translocase TOMM40 and the DNA excision repair protein ERCC1, and it is specifically altered in Opc; suggesting potential additional roles of oligodendrocyte lineage cells mediating AD genetic risk.

Overall, these results uncover stage and cell type specific transcriptional responses to AD pathology, suggesting early and strong activation of metabolic, cell cycle, and immune alterations in specific cell types; along with an incremental activation of neurotransmission and apoptotic processes involving primarily neuronal cells.

### Stage-dependent transcriptional alterations

We next focused on alterations with contrasting and stage-dependent responses to AD pathology, by directly comparing late and early module association changes (**Fig. 5a**). Astrocytes showed the most extreme stage-dependent responses, suggesting that these cells are dynamically responsive to AD pathology in stage progression. We observed increased early-stage expression of genes involved in cholesterol metabolism and transport (e.g., LDLR, SOAT1, ABCA1, HMGCS1, MSMO1), and in inflammation (e.g., IL6R, JAK2, STAT1) (both captured by module M4). Inflammation genes were also increased in OPCs, albeit less strongly. Astrocytes in addition showed early-stage decreased expression of neurotransmission and fatty acid metabolism genes, including glutamate transporters, enzymes, and receptors (e.g., SLC1A2, GLUD1, GLS, GRIA2; module M6); suggesting dysregulation in homeostatic support for neuronal neurotransmission. Finally, astrocytes showed increased late-stage expression of β-amyloid toxicity and neuroinflammation genes (e.g., CSF1, CD44, FOS, IKBKB; module M5), both processes commonly reported as potentially having mediating roles in AD progression^37,38^ (**Fig. 4e, f; Fig. 5a**).

**Figure 5.**
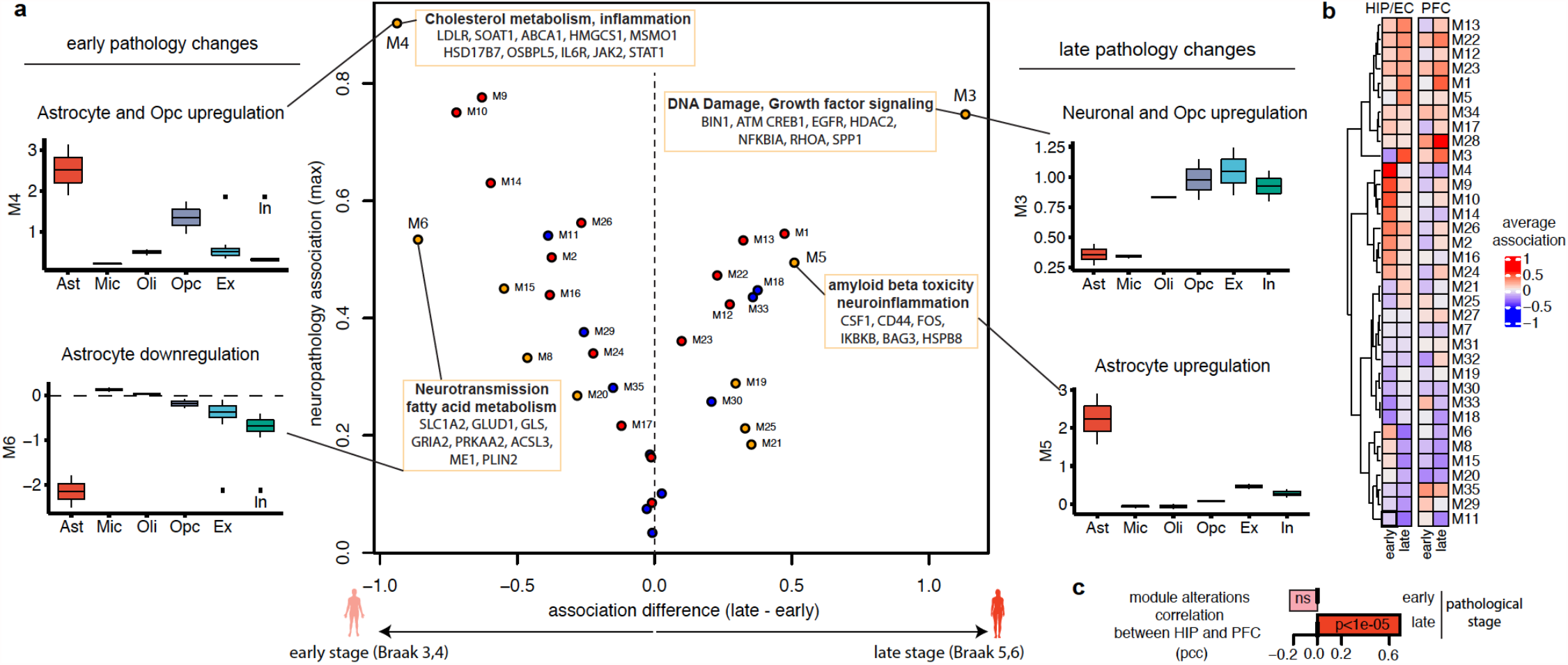
Stage-dependent expression alterations and cross regional convergence. **a**, Module association scores ordered by the difference in the magnitude of association with late and early neuropathology stages (x-axis) and the maximum absolute association score (y-axis). Modules that capture genes with consistent downregulation across stages as highlighted in blue, those with consistent upregulation with red, and those in inconsistent changes in orange. Representative molecular pathways and genes are highlighted for the modules with most extreme stage-dependency (rectangles), along with their association scores across cell types (boxplot). **b**, Average module association scores by pathology stage (early, Braak 3 or 4; late Braak 5 or 6) and brain region (HIP/EC, left; PFC, right). Association scores are computed by the negative logarithm of the p-values estimated by a negative binomial mixed model signed by the directionality of expression change. **c**, Correlation between HIP/EC and PFC average module association in early and late pathology stages.

In addition to astrocytic responses, we also detected strong but less specific stage-dependent alterations in neurons. In particular, module M3 captured late-stage increased expression of genes involved in DNA damage, growth factor signaling, and cellular stress (e.g., BIN1, CREB1, HDAC2, EGFR, NFKBIA, RHOA, SPP1, SESN2, BAG6, CDKN2AIP, FZR1). This response primarily occurs in excitatory neurons, but also in oligodendrocyte lineage cells and inhibitory neurons, suggesting broad stress response activation in late pathology stages.

To investigate the extent to which the molecular pathological signatures recovered herein in early affected regions extend to those affected late in AD progression, we compared our results with data from the prefrontal cortex. To this end, we first reanalyzed our previously published single-nucleus transcriptomes of the prefrontal cortex^13^ to infer analogous AD-associated expression differences and association scores in the two pathological stages used here (early/limbic Braak 3 or 4, and late/neocortical Braak 5 or 6); while accounting for age, sex, and postmortem time intervals (**Supplementary Table S7)**. We then used prefrontal cortex data to quantify the average stage-dependent association of the module alterations uncovered across cells of the hippocampal and entorhinal areas (**Fig. 5b**). We found that only late-stage module responses were consistently recovered in both regions (cor=0.69, p-val=4.5e-06), while early responses diverged (**Fig. 5c**); highlighting the importance of sampling tissue from early-affected regions to capture the unique cellular environment of early molecular pathology, and suggesting that late-stage cellular responses to a certain degree tend to converge towards multicellular molecular signatures suggestive of neurotransmission dysregulation, cellular stress, and DNA damage.

### Specificity of neuronal molecular pathology

We next sought to pinpoint specific anatomical structures and neuronal subtypes where AD-associated changes preferentially occur, by quantifying the association between gene expression levels and neurofibrillary tangle pathology (NFT) burden for each neuronal subtype separately (**Supplementary Table S9**). Given the intracellular and selective character of NFT pathology ^22^, we used a targeted approach to uncover molecular processes potentially mediating neuronal dysfunction, specifically focusing on anatomical structures and neuronal populations that are robustly annotated, directly targeted during tissue recollection, and expected to include neurons that are vulnerable to neurodegeneration^21,39^. For glutamatergic neurons, we focused on CA1, CA3, and DG subtypes from the hippocampus (**Fig. 6a**) and ECL2, ECL2/3, ECL3, ECL5, and ECL6 from the entorhinal cortex (**Fig. 6b**).

**Figure 6.**
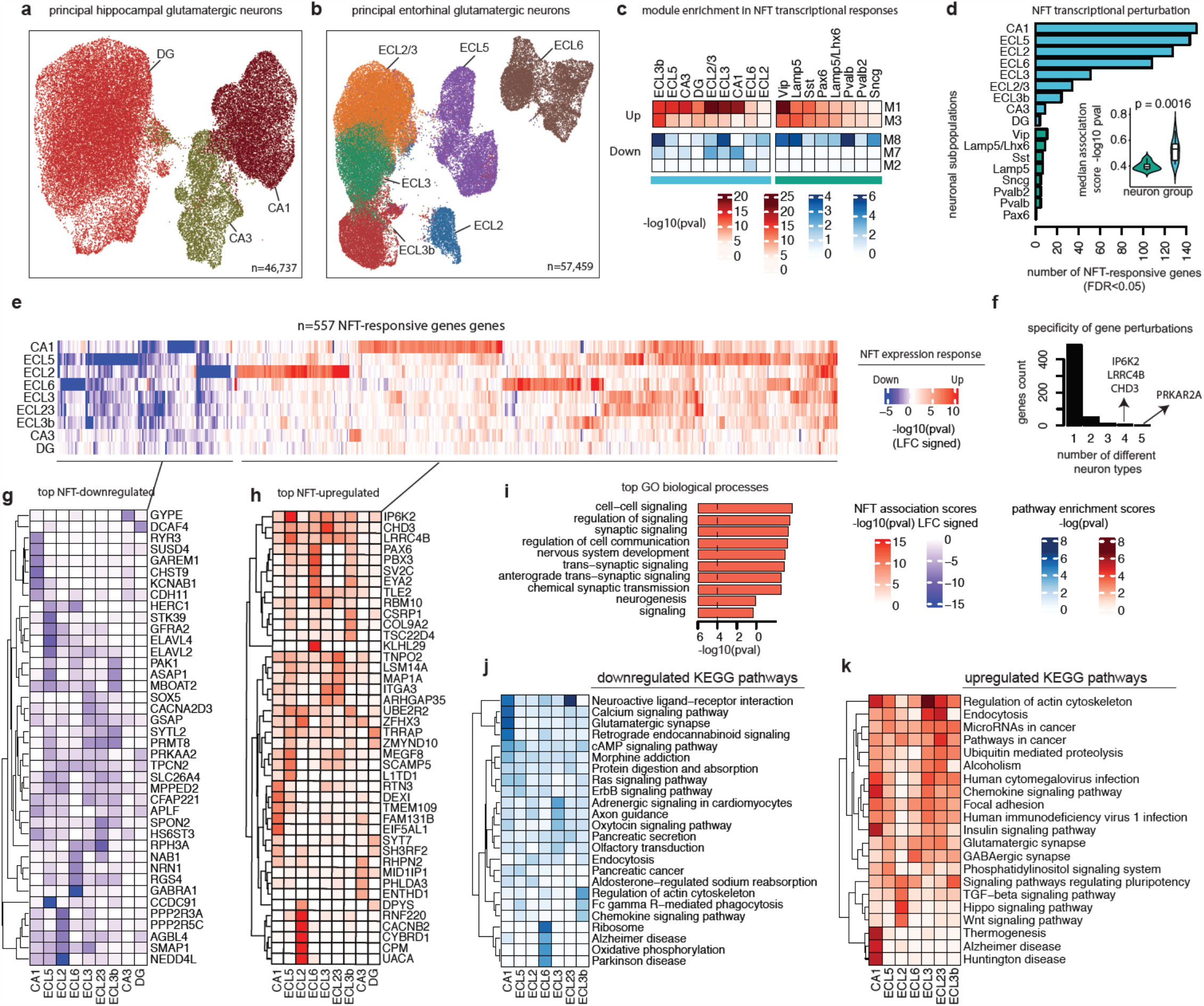
Neuron-type specific molecular pathology. **a-b**, Two-dimensional ACTIONet representation of principal hippocampal glutamatergic neurons (n=46,737) (**a**) and principal hippocampal glutamatergic neurons (57,459) (**b**) labelled by subpopulations deduced from integrative data interpretation. **c**, Enrichment test assessing and quantifying whether module genes tend to present NFT-responses within neuron types. **d**, NFT-responsive gene count across neuron types. **e**, Association patterns of significant (FDR<0.05, NBMM) NFT-responsive genes across neuron types. **f**, Number of genes perturbed in one or more neuron types in response to NFT. **g-h**, Top 5 downregulated (**g**) and upregulated (**h**) genes in response to NFT within neuron types. **i**, Top GO biological processes overrepresented within NFT-responsive genes. **j-k**, Top 3 downregulated (**j**) and upregulated (**k**) KEGG pathways in response to NFT within neuron types.

We first investigated whether the AD-associated gene modules previously discovered at cell-type resolution capture transcriptional NFT responses occurring broadly or preferentially across anatomical structures and neuronal subtypes. We found that several of these modules overlap with NFT-associated changes occurring preferentially across neuronal subtypes, highlighting the relevance of a targeted subpopulation-level analysis (**Fig. 6c**). Among NFT-upregulated modules, we found that neurotransmission and apoptosis-related module M1 was broadly upregulated across most glutamatergic subpopulations, with stronger changes in CA1 and in superficial to midlayer EC neurons (ECL2-L3) among excitatory neurons, and only in Vip among inhibitory neurons. By contrast, cellular stress and DNA damage-related M3 showed specific up-regulation in ECL3 neurons, but only modest NFT-associated changes in other anatomical regions. Among NFT-downregulated modules, metabolism-related M8 captured changes in specific subsets of both excitatory and inhibitory neurons, suggesting shared dysregulated metabolic changes across neuronal groups, while synaptic signaling and transmission-related M7 captures genes specifically downregulated in CA1 and EC L2/3 glutamatergic neurons.

We next sought to uncover additional alterations that were not captured at cell-type resolution, by analyzing neuronal subtype-specific responses to NFT pathology directly. These responses showed that glutamatergic neuron types are globally more transcriptionally responsive to NFT pathology than GABAergic types, as measured both by a global summary statistic (median absolute association scores -log10(pval), NBMM), and by the total number of genes with a significant response to NFT (FDR<0.05, NBMM) (**Fig. 6d**). Within glutamatergic subpopulations, principal pyramidal cells showed more changes than granule cells of the dentate gyrus, with pyramidal neurons of the CA1 field changing the most, followed by EC neurons of layers L5 and L2, and with CA3 neurons changing the least (**Fig. 6d**).

We identified a total of 557 highly specific NFT-responsive genes across the 9 neuronal subtypes considered (**Fig. 6e-h, Supplementary Table 10**), with the vast majority (N=489, 87.7%) presenting expression changes in only one anatomical region and neuronal subtype and only a few being perturbed in 4 or more anatomical regions and neuronal subtypes (**Fig. 6f**). Among recurrent gene perturbations we found the cAMP-dependent signaling molecule PRKAR2A involved in the regulation of lipid and glucose metabolism, the immune responsive enzyme IP6K2, the cell adhesion molecule LRRC4B involved in excitatory synaptic formation, and the DNA binding protein CHD3 involved in chromatin remodeling; all of which were overexpressed across anatomical regions and neuronal subtypes. Neuronal-subtype specificity is also evidenced in the fact that only a fraction (189, 33.3%) of NFT-responsive genes were also found to be differentially expressed at cell-type resolution and by the molecular processes involved, which include key processes for neuronal function, such as cellular communication, synaptic signaling, and neurotransmission (**Fig. 6i)**.

Consistent with signatures of intercellular proteostatic stress, we observed that NFT-upregulated genes involve actin cytoskeleton and endocytosis primarily in CA1 and ECL2/3 neurons, and ubiquitin mediated proteolysis more broadly. In addition, we detected recurrent upregulation of several viral-related pathways, suggesting an inflammatory neuronal component consistent with recent findings in mouse models and human cortical AD samples^40^. Unlike EC neuron types, NFT-responsive genes in CA1 neurons involve the upregulation of KEEG neurodegenerative disease pathways, including AD, Hungtington’s, and Parkinson’s disease; along with increased insulin and chemokine signaling, and the specific downregulation of glutamatergic synaptic and calcium signaling genes. Together these changes are suggestive of concomitant intercellular stress and neurotransmission dysregulation. EC neurons, in contrast, show partial upregulation of glutamatergic and GABAergic synaptic genes in both superficial (L2/3, L3) and deep layer (L6) neurons, while the later also shows specific downregulation of neurodegenerative disease pathways.

## Discussion

We report the first (to our knowledge) single-nucleus transcriptomic atlas encompassing the aged human hippocampus and entorhinal cortex in the context of early and late stages of AD pathology. Using integrative data analysis, we reproducibly identified brain cell types and anatomically annotated subtypes, including specific glutamatergic and GABAergic neuronal subpopulations largely consistent with mouse and human anatomical and cellular transcriptomic signatures. We show that the anatomical topography of human hippocampal structure is strongly reflected in its single-cell neuronal molecular topography: the closer the hippocampal substructures are anatomically, the more similar their neuronal transcriptomes are. These results are consistent with tissue-level observations in the brain as a whole^32^ and across the neocortex ^31^, and our findings demonstrate that single-cell analysis can recover the cell types and subtypes underlying spatial anatomical associations observed at tissue-level resolution. Our single-cell datasets and computational integrative analyses revealed that the architecture of the human hippocampus transcriptome is characterized largely by structured glutamatergic neuronal heterogeneity and the ratio of neuronal to glial signatures, rather than GABAergic heterogeneity.

Our characterization of early-stage (Braak 3,4) and late-stage (Braak 5,6) transcriptional changes uncovered consistent and coordinated expression changes that suggest a molecular manifestation of pathology that is highly stage- and cell type specific and broadly consistent across hippocampal and entorhinal major cell types. Only late-stage expression changes are globally consistent with those observed in the prefrontal cortex, suggesting convergent late-stage responses across these cortical and subcortical regions, but a distinct and specific cellular response to the local hippocampal environment in early stages of pathological progression. Future work will help elucidate to what extent these observations generalize to other brain regions, and whether such consistency is related with the lower heterogeneity of glial relative to neuronal cells observed across studies^13,17,23,31^.

While most previous work has focused on characterizing dysregulated cell type specific functions that might contribute to AD, including inflammatory responses of microglia^41^ and astrocytes^41,42^, potential myelination defects of affected oligodendrocyte lineage cells^13^, and pericyte-mediated blood-brain barrier dysfunction^43,44^; our data suggest that alterations to the intercellular modulatory role of glial in neurotransmission might have a contributing role in AD. We found extensive expression and recurrent transcriptional alteration of genes directly involved in synaptic communication and transport mechanisms in both astrocytes and oligodendrocyte lineage cells, including glutamate transporters (SLC1AA2), processing enzymes (GLUD1, GLS), and receptor subunits (e.g., GRM3, GRID2, GRM7). Astrocytes showed a particular dynamic and stage-dependent response to AD pathology, involving early dysregulation of cholesterol metabolism and neurotransmission, but late neuroinflammation; a pattern not observed in other glial cells, which our data suggest present more consistent changes in pathology progression.

In neuronal cells, our results show that glutamatergic neurons are more affected transcriptionally than GABAergic neurons by NFT pathology, and that distinct subpopulations vary in their degree of transcriptomic alteration. Patterns of alteration in glutamatergic neurons reveal an anatomical pattern of neuronal impact across human hippocampal substructures, with CA1 pyramidal neurons most affected, followed by pyramidal neurons of entorhinal deep and mid layers, and ending up with CA3 neurons and DG granule cells which show the least effect. These observations are consistent with the selective vulnerability of CA1 and EC pyramidal cells^11,21,39^, and suggest that observed transcriptomic responses may reflect molecular processes relevant for neuronal dysfunction. Potentially among the latter, we found that CA1 neurons show simultaneous downregulation of homeostatic neuronal functions and upregulation of processes suggestive of neuronal stress, including metabolic alterations, cytoskeletal transport dysfunction, inflammatory responses, and genes involved in neurodegeneration. The resulting data resource put forward herein will open up opportunities for future studies following a more targeted approach to human neuronal vulnerability and its molecular basis.

Finally, although rich and extensive, the breadth of cellular diversity discovered in this study might still be limited by lack of samples from anatomical substructures targeted by microdissection. We approached this limitation computationally by including the transcriptomic signatures of existing, anatomically-annotated samples from mouse and human in our analysis. It should be noted that our study was primarily targeted to the hippocampal and entorhinal region, information of the parahippocampal and subicular structures is limited. Future work will complement our data by considering extensive anatomical microdissection of hippocampus-surrounding regions to further characterize the progression of neuronal alterations. Overall, our single-cell analysis of the human hippocampus and entorhinal cortex highlights the diversity of cells within the hippocampal formation and the effect of cellular specificity and pathological stage progression in AD. Our data and results will provide a valuable resource to guide the mechanistic study of neuronal vulnerability in the context of the human brain, and to contrast observations and predictions from animal and human in-vitro models of the disease.

## Methods

### Data reporting

No statistical methods were used to predetermine sample size.

### ROSMAP subject selection

65 individuals were selected from the The Religious Orders Study and Rush Memory and Aging Project (ROSMAP), two longitudinal cohort studies of aging and dementia that collect extensive clinical data and perform detailed post-mortem pathological evaluations, as previously described^28^. For the purpose of this study, we considered a pathological diagnosis of AD to define subject groups. Based on the modified NIA-Reagan diagnosis of Alzheimer’s disease, 31 individuals were classified as having a positive pathological diagnosis of AD (AD group) and 34 individuals a negative diagnosis (no AD pathology group, NoAD). Pathological evaluation relies on both neurofibrillary tangles (Braak) and neuritic plaques (CERAD), and it is done without knowledge of clinical information. Pathology groups were balanced by male:female ratios (16:15 in AD, 17:17 in NoAD), age (72-95 years, average 85:84), and postmortem interval (0.86-12 hours, average 6:6.5). Three additional groups were defined based on involvement of neurofibrillary tangle (NFT) pathology as assessed by Braak staging: reference stage (Braak stage 0, 1, 2; n=31), early/limbic stage (Braak stage 3 or 4, n=17); and late/neocortical stage (Braak 5 or 6, n=17). Detailed description on clinical and pathological data collection has been previously reported^45^. The Religious Orders Study and Rush Memory and Aging Project were approved by an Institutional Review Board (IRB) of Rush University Medical Center. Informed consent, an Anatomical Gift Act, and a repository consent were obtained from each subject. ROSMAP data can be requested at https://www.radc.rush.edu.

### Isolation of nuclei from frozen post-mortem brain tissue

The protocol for the isolation of nuclei from frozen post-mortem brain tissue was adapted from a previous study (Mathys et al. Nature 2019). All procedures were carried out on ice or at 4°C. In brief, post-mortem brain tissue was homogenized in 700 µl homogenization buffer (320 mM sucrose, 5 mM CaCl_2_, 3 mM Mg(CH_3_COO)_2_, 10 mM Tris HCl pH 7.8, 0.1 mM EDTA pH 8.0, 0.1% IGEPAL CA-630, 1 mM β-mercaptoethanol, and 0.4 U µl^−1^ recombinant RNase inhibitor (Clontech)) using a Wheaton Dounce tissue grinder (15 strokes with the loose pestle). Then the homogenized tissue was filtered through a 40-µm cell strainer, mixed with an equal volume of working solution (83% OptiPrep density gradient medium (Sigma-Aldrich), 5 mM CaCl_2_, 3 mM Mg(CH_3_COO)_2_, 10 mM Tris HCl pH 7.8, 0.1 mM EDTA pH 8.0, and 1 mM β-mercaptoethanol) and loaded on top of an OptiPrep density gradient (750 µl 30% OptiPrep solution (30% OptiPrep density gradient medium,134 mM sucrose, 5 mM CaCl_2_, 3 mM Mg(CH_3_COO)_2_, 10 mM Tris HCl pH 7.8, 0.1 mM EDTA pH 8.0, 1 mM β-mercaptoethanol, 0.04% IGEPAL CA-630, and 0.17 U µl^−1^ recombinant RNase inhibitor) on top of 300 µl 40% OptiPrep solution (40% OptiPrep density gradient medium, 96 mM sucrose, 5 mM CaCl_2_, 3 mM Mg(CH_3_COO)_2_, 10 mM Tris HCl pH 7.8, 0.1 mM EDTA pH 8.0, 1 mM β-mercaptoethanol, 0.03% IGEPAL CA-630, and 0.12 U µl^−1^ recombinant RNase inhibitor). The nuclei were separated by centrifugation (5 min, 10,000 g, 4 °C). A total of 100 µl of nuclei was collected from the 30%/40% interphase and washed with 1 ml of PBS containing 0.04% BSA. The nuclei were centrifuged at 300*g* for 3 min (4 °C) and washed with 1 ml of PBS containing 0.04% BSA. Then the nuclei were centrifuged at 300*g* for 3 min (4 °C) and re-suspended in 100 µl PBS containing 0.04% BSA. The nuclei were counted and diluted to a concentration of 1,000 nuclei per microliter in PBS containing 0.04% BSA.

### Droplet-based snRNA-seq

For droplet-based snRNA-seq, libraries were prepared using the Chromium Single Cell 3′ Reagent Kits v3 according to the manufacturer’s protocol (10x Genomics). The generated snRNA-seq libraries were sequenced using NextSeq 500/550 High Output v2 kits (150 cycles) or NovaSeq 6000 S2 Reagent Kits.

### snRNA-seq data preprocessing

Gene counts were obtained by aligning reads to the GRCh38 genome using Cell Ranger software (v.3.0.2) (10x Genomics). To account for unspliced nuclear transcripts, reads mapping to pre-mRNA were counted. After quantification of pre-mRNA using the Cell Ranger count pipeline, the Cell Ranger aggr pipeline was used to aggregate all libraries (without equalizing the read depth between groups) to generate a gene-count matrix. The Cell Ranger 3.0 default parameters were used to call cell barcodes.

### Cell inclusion criteria

Outlier cells with less than 500 or more than 10,000 genes detected were excluded, and only genes detected in at least 10 cells were considered. The following quality measures were quantified for each cell: (1) the number of genes for which at least one read was mapped (indicative of library complexity); (2) the total number of counts; (3) the total number of counts mapping to mitochondrial genes, and (4) the percentage of reads mapped to mitochondrial genes (used to approximate the relative amount of endogenous RNA and commonly used as a measure of cell quality). Outlier cells with extremely high total counts and/or mitochondrial read percentage were excluded. Cut-off values for exclusion were determined empirically through examination of descriptive plots of quality metrics. Initial QC metric computation and filtering was performed using the software package scanpy^46^. Nuclear-encoded protein coding genes were considered for downstream analyses.

### Clustering analysis and QC filtering

All clustering analyses, dimensionality reduction, and visualization steps were performed using our computational analysis framework ACTIONet reported in ^47^ and available at (https://github.com/shmohammadi86/ACTIONet/tree/R-release). Briefly, for each round of clustering, single value decomposition (SVD) is initially performed for feature (gene) dimensionality reduction and selection, producing a low-rank approximation of the normalized count matrix. This reduced data representation is subsequently decomposed multiple times to define a multiresolution and low dimensional cell state representation for each individual cell. This final cell state representation is used to build a cell network or manifold, whose structure captures relationships in transcriptomic state similarity at single-cell resolution. Cell similarity and network construction based on these representations has been shown to recover biological cellular associations with improved performance relative to more conventional methods^47^. To avoid biased cell mixing due to independent sequencing batches or potential technical artifacts, a batch correction step considering subject membership as indicator vector was performed as part of the initial dimensionality reduction. This step is achieved using ACTIONet’s function *reduce*.*and*.*batch*.*correct*.*ace*.*Harmony*, which internally uses the data integration procedure reported in^48^ and implemented in the software package Harmony (https://github.com/immunogenomics/harmony). To identify discrete groups of cells with similar transcriptomic signatures (cell clusters), the Leiden graph-based clustering algorithm^49^ is applied to the resulting cell network. The same methodology was applied in the second clustering round, using cell type annotated subsets of cells as input. As part of QC filtering steps, clusters representing only cells from one individual, suspected to recover doublet cells, or to be composed of presumed low-quality cells, were excluded from downstream analyses. Doublet or low-quality cluster status was determined empirically based on the examination of cell associations in 2D plots, the presence of mixed gene markers from distinct cell types, and extreme QC metric values. The latter relative to those commonly observed in other subclusters of the same cell type. To ensure reproducibility of annotations, and to facilitate high-resolution identification of region-specific and low-abundant cell types, as well as to enforce a more stringent QC filtering process; HIP and EC datasets were analyzed and annotated separately, and a joint manifold was subsequently reconstructed for joint 2D visualization (Extended data Fig. 1c, d).

### Neuron joint clustering analysis

Gutamatergic excitatory and GABAergic inhibitory neurons previously annotated at group level during clustering analyses were extracted separately from the HIP and EC datasets. Cells from both HIP and EC were then analyzed jointly by integrative clustering analysis using ACTIONet’s pipeline, including a Harmony-based batch correction and data integration step. Briefly, normalization, dimensionality reduction, network construction, and clustering steps were applied jointly to neuronal cells from both brain regions, considering subject membership as indicator vector during the reduction-batch correction step. This procedure was applied to Gutamatergic and GABAergic neurons separately and resulted in the identification of the neuronal subpopulations interpreted and annotated throughout the study (Fig. 2a -c, Extended data Fig. 2a, b).

### External data sources

Reference cell type marker genes were obtained from PsychENCODE^29^ and PanglaoDB^30^. Spatial transcriptomic data from mouse hippocampal and parahippocampal regions was obtained from^33^. Neuron-sorted transcriptomic data from mouse principal hippocampal cells was obtained from^34^. Mouse hippocampal and retrohippocampal single-cell transcriptomic data reported in^50^ was obtained from the Allen Institute’s cell type database(https://portal.brain-map.org/atlases-and-data/rnaseq/mouse-whole-cortex-and-hippocampus-smart-seq). Human tissue-resolution transcriptomic data from laser-microdissected hippocampal subregions reported in ^32^ was obtained from^51^. Summary profiles corresponding to all these data sets and used in integrative analyses throughout the study are reported in Supplementary Table S1. Genes genetically associated with AD based on a genome-wide association study (GWAS) were obtained from^10^, considering genes reported as significantly associated with AD in gene-based association tests (GWGAS). MAGMA gene-level GWAS association scores for AD (ID 4091), Schizophrenia (ID 3982) and Type 2 Diabetes (ID 4176) were obtained from (https://atlas.ctglab.nl/traitDB), selecting in each case the most recent study with the largest sample size. Genes associated with neurodegenerative disorders were extracted from genesets curated in the DisGeNET collection^52^, using the keywords *Parkinson, Amyotrophic, Dementia, dementia*, and *Hungtinton* to keep those associated with related degenerative disease. For gene module and pathology associated gene interpretation, the following reference gene sets were extracted from the gene set enrichment analysis server Enrichr^53^: *GO_Biological_Process_2018, KEGG_2019_Human, and Elsevier_Pathway_Collection*. Genes associated with neuropathology and neuroinflammation were extracted from nanoString gene panels (https://www.nanostring.com/).

### Mouse single-cell transcriptomic reanalysis

Cells isolated from hippocampal and parahippocampal areas (hippocampus, entorhinal cortex, para-, post-, and pre-subiculum, and subiculum) annotated with region labels HIP, ENTm, ENTl, PAR-POST-PRE, and SUB-Pro were extracted from the original data downloaded from (https://portal.brain-map.org/atlases-and-data/rnaseq/mouse-whole-cortex-and-hippocampus-smart-seq). Due to its high similarity with one of the human subclusters uncovered herein, the mouse cluster 1_CR was also extracted and included in reanalysis. Subsequently, cells with class label Glutamatergic or GABAergic were separately extracted and reanalyzed. The resulting datasets were normalized, analyzed, and visualized following a similar pipeline to the ones used for human data analysis: the standard pipeline of the ACTIONet framework. To directly contrast human and mouse relative neuron type similarity and localization, reported mouse subclass-level annotations were considered.

### Cell type interpretation and annotation

A transcriptional profile and a transcriptional signature were defined for each cluster, subcluster, cell type, and neuron subpopulations, and these were used to aid interpretation of cell groups and subgroups based on gene expression patterns. A profile was defined by the average expression across group member cells. A signature was defined by the total sum of the pair-wise expression fold-change of a given profile relative to each of the other profiles. Genes with high signature values for a given group represent preferentially expressed genes and tend to correspond to well-known cell type markers. Signatures were used throughout the study to unbiasedly aid interpretation by quantifying the degree to which genes with known biological meaning show unexpectedly high preferential expression values within a given cell group, and to assess global transcriptomic similarity across datasets. Marker genes previously curated as part of the PsychENCODE consortium^29^ were used to assign and quantify cell type interpretations to the different cell groups. Interpretations were independently verified using an additional curated cell-type marker compendium available in PanglaoDB^30^. To compare the cell groups with external transcriptomic data sources, transcriptional profiles and signatures were similarly computed for the external reference dataset. The Pearson correlation coefficients between cell groups and reference dataset signatures were used as measures of similarity/consistency, and top-ranking reference conditions across datasets were examined to determine best-matching and independently supported interpretations and labels (Extended data Fig. 3).

### Preferential gene expression in neuronal subpopulation

Preferential expression was determined by testing whether a gene is overexpressed in cells of a given neuron subpopulation relative to the rest of neurons of the same type (Glutamatergic or GABAergic), while considering cell-subject membership associations. All comparisons were formalized by fitting a negative binomial mixed model to account for cell-subject membership associations. All models were implemented using the R package Nebula^54^. The analysis was conducted in the joint data set, including all annotated neuronal cells from the HIP and EC samples. Genes were considered as preferentially expressed according to the following criteria: FDR < 0.01 and logFC>1.

### Gene expression changes in AD pathology

Gene expression associations were assessed using a fast implementation of a negative binomial mixed model that accounts for both subject-level and cell-level overdispersion^54^. To estimate the association between gene expression and Braak stage groups, while accounting for sex, age of death, postmortem interval. and additional covariates related with cell quality, the following model formulation was used

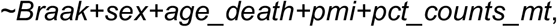

where Braak and sex are discrete variables specifying Braak group (early, Braak 3 or 4; late, Braak 5 or 6; and reference, Braak 0, 1 or 2); and sex, and *age_death, pmi, and pct_counts_mt* are numerical variables specifying the age of death, the postmortem time interval, and the percentage of counts mapping to mitochondrial genes, respectively. Additionally, total counts per cell were used as offset in the model specification. Using this formulation, expression levels in early and late stages were each compared versus the reference stage with no or with low pathology. All association models were implemented using the R package Nebula (https://github.com/lhe17/nebula).

### Neuropathology gene module analysis

For each gene, a gene neuropathology association profile was defined by concatenating gene association scores for each combination of Braak stage (early or late), brain region (hippocampus, entorhinal cortex), and cell type (excitatory neuron, inhibitory neuron, astrocyte, oligodendrocyte, Opc, microglia). Association scores were measured by the logarithm of the p-value estimated by the NBMM multiplied by 1 or -1 when the expression of the gene increased or decreased relative to the reference stage. This resulted in the operational definition of a gene neuropathology association profile as a signed numeric vector, where each of 20 scores measures the degree to which the expression of a gene is positively (overexpression) or negatively (underexpression) associated with a given pathological stage in a given brain region and cell type. To measure gene neuropathology association profile similarity, profiles were first mean-centered and scaled and then used to compute pairwise Pearson correlation coefficients, resulting in a profile similarity matrix (Extended data Fig. 4). To identify groups of genes with consistent profiles (gene modules), while considering the directionality (sing) of associations, the signed version of the Leiden graph-clustering algorithm^49^ as implemented in the function *Leiden*.*clustering* of the ACTIONet package was used. The output of this methodology are groups of genes with measurably similar neuropathological stage associations across brain regions and cell types. Only genes having an association p-value < 0.01 in at least one of the comparisons were considered. To minimize potential false-positive association, only associations of genes whose expression was reliably detected in a given cell type were considered (nonzero read counts in at least 10% of the cells of a given type).

### Neuronal type specific NFT association analysis

Associations between gene expression levels and NFT burden within neuronal subpopulations were assessed using a negative binomial mixed model. To estimate associations, while accounting for sex, age of death, postmortem interval and additional covariates related with cell quality, the following model formulation was used

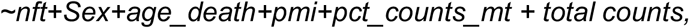

where *nft* represents the measurement of neurofibrillary tangle burden, as determined by microscopic examination of silver-stained slides. Only genes having an estimated FDR<0.05 were considered as NFT-responsive. All association models were implemented using the R package Nebula^54^.

### Gene overrepresentation analysis

Overrepresentation of marker gene sets within cell groups for cell annotation was assessed by permutation tests as implemented in the ACTIONet package function *annotate*.*profile*.*using*.*markers*, considering cell type or cell group signature profiles as input profiles. Briefly, tests assess whether genes as a group tend to have unexpectedly high values, as defined in the input profiles and measured by a z-score statistic. Rank-based gene set enrichment analyses (GSEA) were performed using the R package (https://bioconductor.org/packages/release/bioc/html/fgsea.html). Gene Ontology gene set enrichment analyses were performed using the R package *gprofiler2*^*55*^ considering gene lists as input and the default gene set background.

## Acknowledgement

We acknowledge members of the Tsai and Kellis groups for helpful discussions. This work is partially supported by RF1 AG054012 (to L.-H.T. and M.K.), RF1 AG062377 (to L.-H.T. and M.K.), CureAlzheimer’s Fund (to L.-H.T. and M.K.).

## Author contributions

J.D.-V., D.A.B., M.K., and L.-H.T. conceived and designed the study. M.K. and L.-H.T. directed and coordinated the study. J.D.-V., and S.M developed and implemented the computational framework, and performed data analysis. H.M., B.R., H.J., and A.N. performed profiling experiments. J.D.-V. wrote the manuscript with input from all authors.

## Declaration of interests

The authors declare no competing interests.

## Extended Data Figure legends

**Extended Data Fig. 1.**
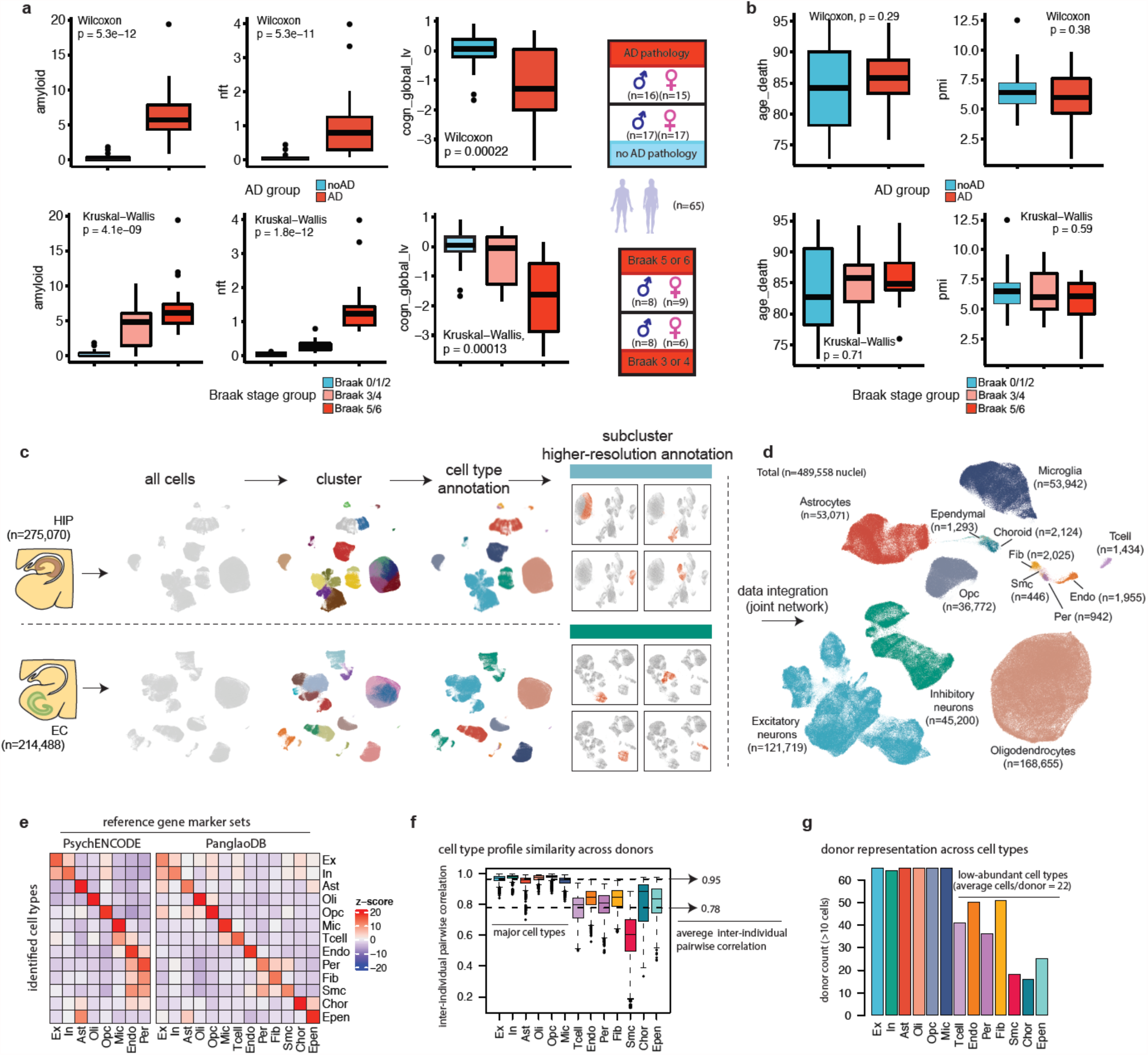
Sample pathological status and cell-type characterization. **a**, Comparison of β-amyloid (left) neurofibrillary tangle (middle) and global cognition (right) measurements between AD groups. **b**, Comparison of age and postmortem time interval (pmi) between AD groups. P-values based on two-sided wilcoxon rank-sum test for AD vs no AD comparisons, and Kruskal–Wallis test for differences across Braak stages. **c**, Clustering analysis workflow. **d**, Two-dimensional ACTIONet projection of all annotated cells across hippocampal and entorhinal samples (n = 489,558 from 31 pathology and 34 no-pathology individuals). **e**, Enrichment analysis (resampling test) within each of the annotated cell types (rows) of genes previously identified as markers by two independent resources (columns). **f**, Interindividual pairwise profile correlation values (Pearson correlation coefficients) by cell type (columns). **g**, Number of individuals for which at least 10 cells of a given type were recovered.

**Extended Data Fig. 2.**
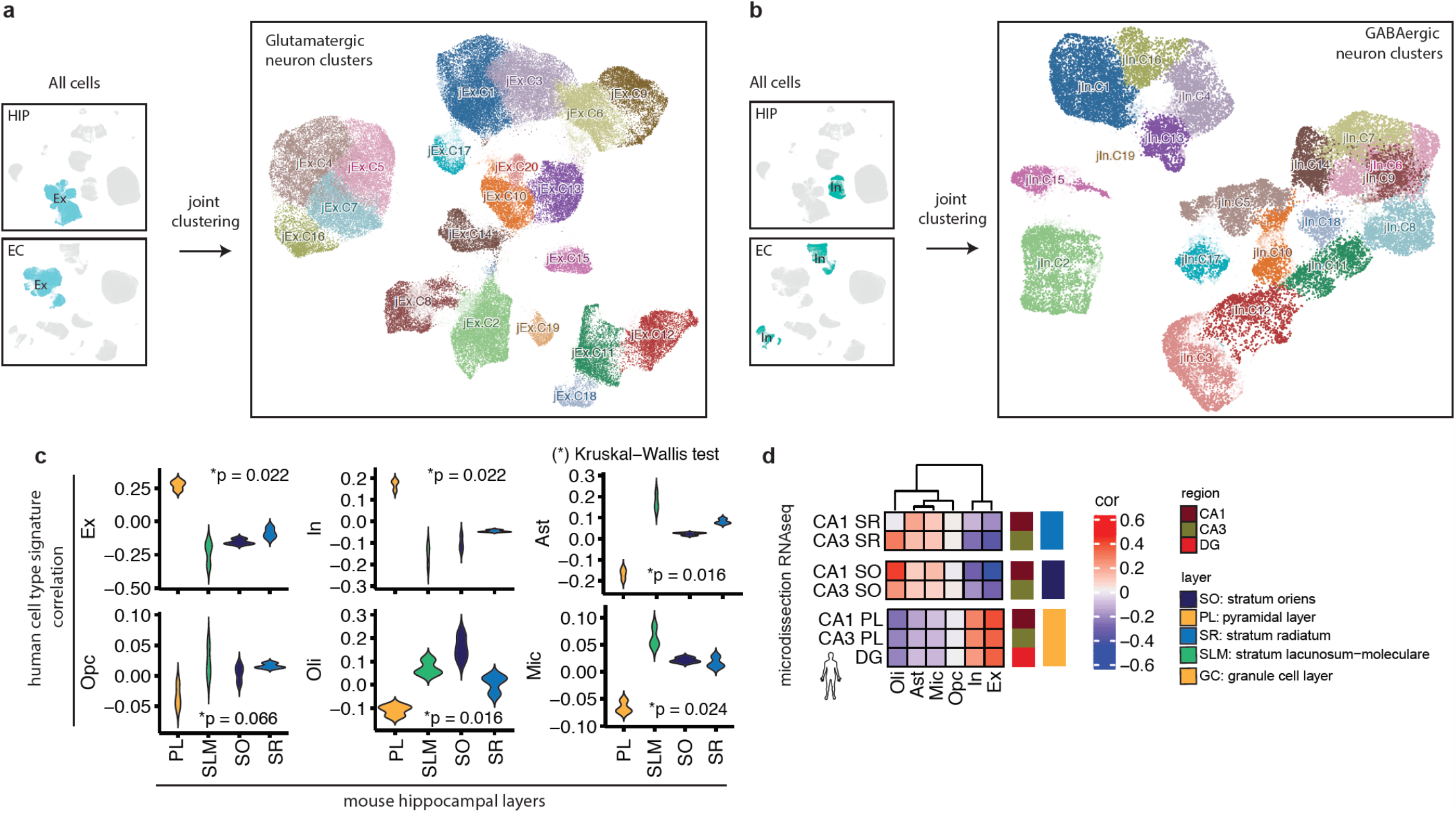
Joint neuronal clustering and hippocampal cytoarchitectural interpretation. **a-b**, Joint clustering analysis of HIP and EC neurons for glutamatergic (**a**) and GABAergic (**b**) groups labeled by cluster in Two-dimensional ACTIONet cell representations. **c**, Comparison of correlation scores between each human cell type signatures and signatures of hippocampal layers computed from mouse spatial transcriptomics data. Comparisons are performed across layers using a Kruskal–Wallis test. **d**, Similarity between human cell type profiles and human hippocampal layered signatures obtained by laser capture microdissection followed by RNA sequencing (LCM-seq) from human hippocampal tissue.

**Extended Data Fig. 3.**
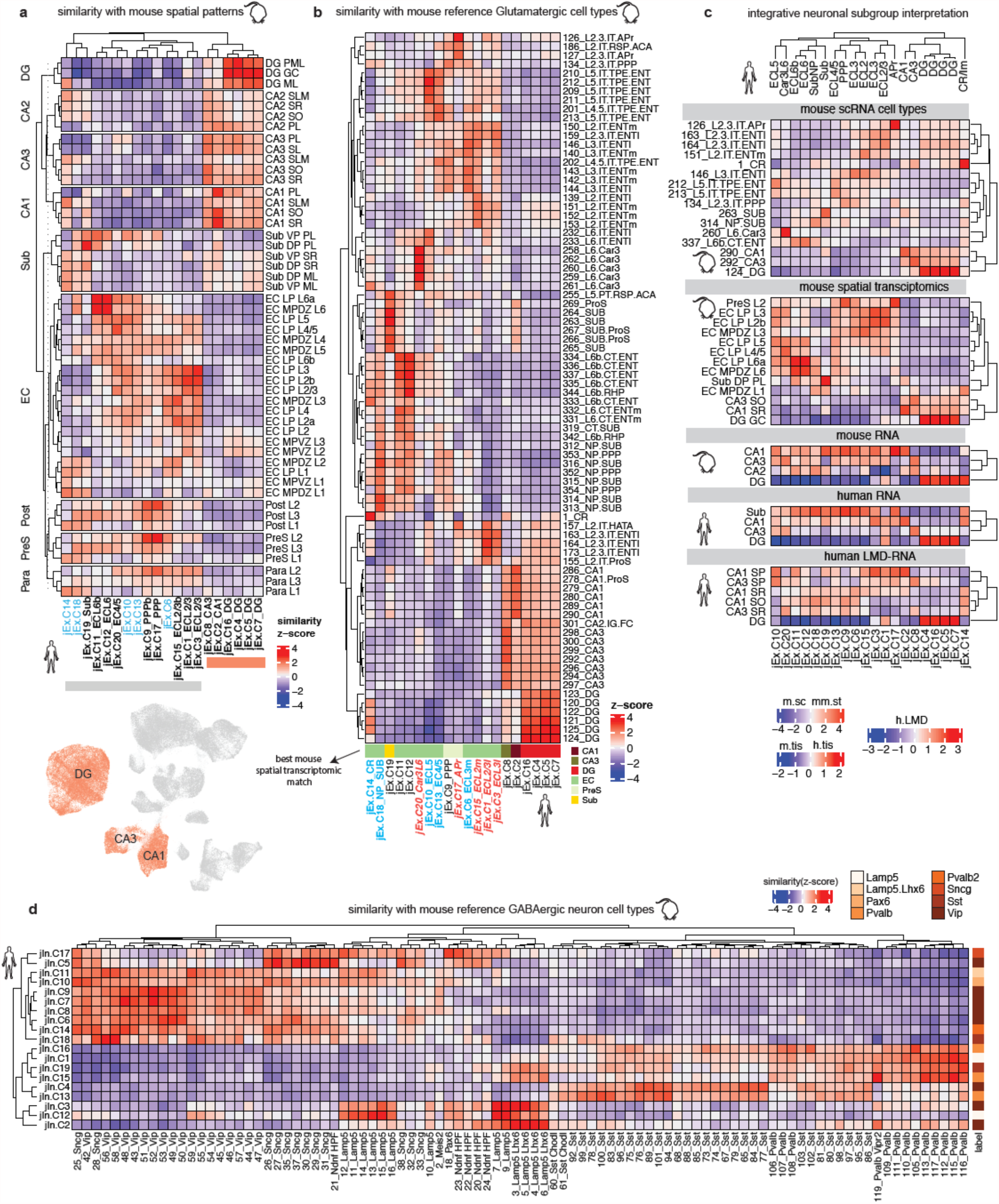
Integrative data analysis for human neuronal interpretation. **a-b**, Similarity between human glutamatergic neuronal subpopulation signatures (columns) and all mouse hippocampal and retrohippocampal substructure signatures computed from spatial transcriptomics data (rows) (**a**), and all mouse hippocampal and retrohippocampal reference transcriptional cell types reported by the Allen Institute (rows) (**b**). **c**, Integrative analysis of similarity measures for human glutamatergic neuronal subpopulations versus mouse signatures at single-cell, spatial, and cell type resolution (top three heatmaps, respectively), and versus human tissue and layer resolution (bottom two heatmaps, respectively). **d**, Similarity between human GABAergic neuronal subpopulation signatures (columns) and all mouse hippocampal and retrohippocampal reference transcriptional GABAergic cell types reported by the Allen Institute (rows).

**Extended Data Fig. 4.**
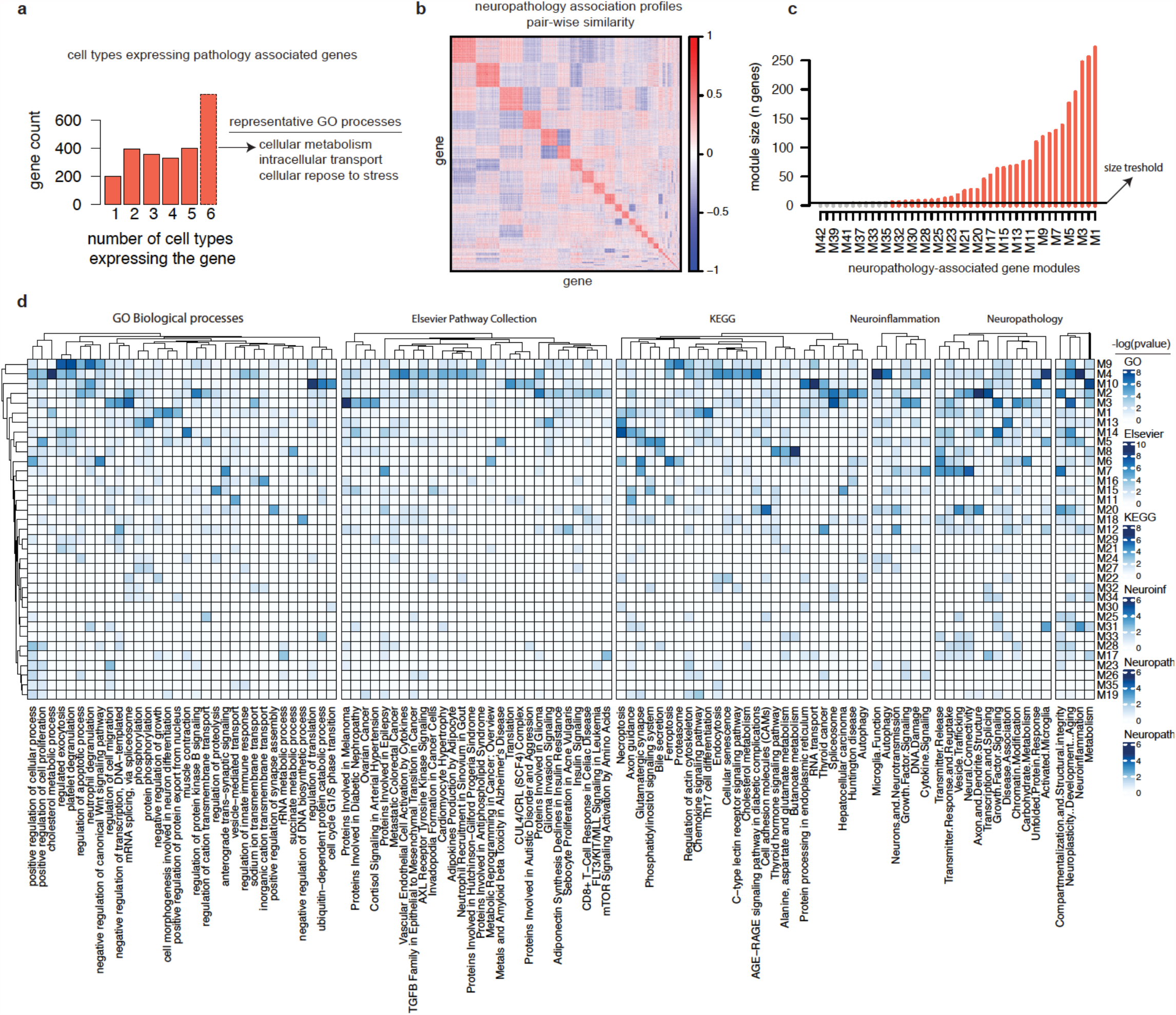
Module analysis of neuropathology associated genes. **a**, Number of genes expressed at base label (nonzero counts in at least 10% of the cell of a given cell type) in one or more cell types. **b**, Similarity between pairs of 2,495 neuropathology association profile. **c**, Module size all identified modules measured in number of genes. Red lines highlight modules passing the arbitrary cut-off of 5 genes. **d**, Enrichment scores of molecular pathways and processes (columns) from multiple references within modules (rows). Enrichment was measured by testing overrepresentation based on expectations from hypergeometric distribution.

## Supplementary tables

**Supplementary Table S1. Human and mouse reference expression profiles**

**Supplementary Table S2. Human cell type and neuronal subpopulation expression profiles Supplementary Table S3. Neuronal subpopulation comparison and interpretation summary Supplementary Table S4. Neuronal subpopulation preferentially expressed genes Supplementary Table S5. Region, cell type, and AD trait specific expression changes**

**Supplementary Table S6. Neuropathology-associated genes cell type expression patterns and GO categories**.

**Supplementary Table S7. Gene neuropathology association profiles. Supplementary Table S8. Gene Module annotation and interpretations**

**Supplementary Table S9. Association between gene expression levels and the burden of NFT pathology across neuronal types**.

**Supplementary Table S10. NFT-responsive genes**

## Notes

### Competing Interest Statement

The authors have declared no competing interest.

